# Supplementation of a lacto-fermented rapeseed-seaweed blend promotes gut microbial- and gut immune-modulation in weaner piglets

**DOI:** 10.1101/2020.09.22.308106

**Authors:** Yan Hui, Paulina Tamez-Hidalgo, Tomasz Cieplak, Gizaw Dabessa Satessa, Witold Kot, Søren Kjærulff Søren, Mette Olaf Nielsen, Dennis Sandris Nielsen, Lukasz Krych

## Abstract

The direct use of medical zinc oxide (ZnO) in feed will be abandoned after 2022 in Europe, leaving an urgent need for substitutes to prevent post-weaning disorders. This study assessed whether rapeseed meal added two brown macroalagae species (*Saccharina latissima* and *Ascophylum nodosum*) and fermented using lactic acid bacteria (FRS) could improve piglet performance and gut health. The weaned piglets were fed one of three different feeding regimens (n = 230 each): basal diet, 2.5% and 5% FRS from day 28 of life to day 85. The piglets fed with 2.5% FRS presented superior phenotype with alleviated intraepithelial and stromal lymphocytes infiltration in the gut, enhanced colon mucosa barrier as well as numerically improvements of final body weight. Colon microbiota composition was determined using amplicon sequencing of the V3 and V1 – V8 region of the 16S rRNA gene using Illumina Nextseq and Oxford Nanopore MinION sequencing, respectively. The two amplicon sequencing strategies showed high consistence between the detected bacteria. Both sequencing technologies showed that the FRS fed piglets had a distinctly different microbial composition relative to the basal diet. Compared with piglets fed the basal diet, *Prevotella stercorea* was verified by both technologies to be more abundant in the FRS piglets, and positively correlated with colon mucosa thickness and negatively correlated with blood levels of leucocytes and IgG. In conclusion, FRS supplementation improved gut health of weaner piglets, and altered their gut microbiota composition. Increasing the dietary inclusion of FRS from 2.5% to 5% did not cause further improvements.

## Introduction

A healthy hindgut is essential for the optimal nutrient utilization and health of the host. Essentially, bacteria largely ferment and digest macronutrients such as insoluble fibers and undigested proteins in the colon (Zeng 2014) (Jha and Berrocoso 2016). Bacterial metabolism benefits colonic cells by providing butyrate and the immune system by providing acetate and propionate (Zeng 2014). However, the process of bacterial colonization, which brings benefits for the host, also brings the risk of infection-inflammation responses. Thus, bacterial association and communication with the immune system balances on a thin line for preserving homeostasis (Peterson and Artis 2014) between the learning of what to attack and what to tolerate (Scudellari 2017).

In pig production, the weaning period is characterized by a change in diet from milk to solid feed, separation from the mother and aggregation in a pen with piglets from other litters. It is the most stressful period in a pig’s life and often lead to morbidities like diarrhea. In-feed zinc-oxide used to be a prevalent choice for prophylaxis, but according to the European Union regulations, zinc-oxide shall no longer be directly used in feed or water from 2022 (Satessa et al. 2020a). This leaves an urgent need for new prophylactic substitutes. Additive inclusion of pre-fermented feeds has been regarded as a promising strategy to ameliorate these post-weaning disorder issues for its effective act to improve gastrointestinal health and enhance livestock performance in production (Hu et al. 2008) (Heres et al. 2003) (Wang et al. 2018). The process of microbial fermentation degrades antinutritional compounds and macronutrients in feed, which increases the nutrient bioavailability and nutritional value (Canibe et al. 2007) (Mukherjee et al. 2015). Besides, the dominant microorganisms in fermented feed have been proposed to inhibit the overgrowth of opportunistic pathogens thus sustaining gut microbiome balance (Plumed-Ferrer and von Wright 2009) and thereby promote immune functions of the host (Zhou et al. 2015). A meta-analysis has shown that fermented feed is able to boost the growth and performance of both weaner and growing pigs (Xu et al. 2020). Supplementation of fermented rapeseed meal also improves the intestinal morphology of broiler chickens (Hu et al. 2016).

Rapeseed meal is a by-product after the oil has been extracted and is generally used as a protein source in animal diets (van der Spiegel et al. 2013). However, it has a lower protein digestibility as compared to soybean meal. Brown seaweed is acknowledged as a good source of health-promoting phytochemicals (Cherry et al. 2019) with bioactive compounds like laminarin (Zargarzadeh et al. 2020), but it has poor digestibility as well. The fermentation with lactic acid bacteria of a rapeseed-seaweed (FRS) blend could improve the health and nutritional value by reducing the naturally present anti-nutritional factors and by hydrolyzing protein and fibers down to a more soluble matrix. We have previously reported that dietary supplementation with FRS could contribute favorably in the development of a healthy functional gut barrier as well as the production performance of piglets in the weaning period (Satessa et al. 2020a).

In the present study, we tested whether addition of FRS to weaner diets will affect modulation of the gut microbiome and the immune function. We chose weaning as the physiological stress situation to investigate this process and to learn how dietary induced gut modulation translates into body weight gain and gut health or illness later in life. To study gut microbial composition in response to dietary intervention, we have used two sequencing strategies of 16S rRNA gene amplicons and compared their performance: 2nd generation (Illumina, NextSeq) and 3rd generation (Oxford Nanopore Technologies, MinION).

## Materials and Methods

### Preparation of fermented rapeseed meal-seaweed feed

The fermented rapeseed meal-seaweed (FRS) feed was provided by FermentationExperts (Denmark), which was a blend of rapeseed meal (*Brassica napus*), wheat bran (*Triticum eastivum*) and two types of brown seaweed (*Saccharina latissima* and *Ascophylum nodosum*) prepared via a controlled 2-step solid state fermentation. The inoculum consists of three lactic acid bacteria: *Pediococcus acidilactici* (DSM 16243), *Pediococcus pentosaceus* (DSM 12834) and *Lactobacillus plantarum* (DSM 12837). The addition of the inoculant controlled the process by acidifying the blend within the first 24 hours, and assuring an almost entirely anaerobic process. The process continued for 11 days at 38 °C. The fermented material was then dried in a spin flash dryer, with a temperature setting and pass-through-speed that preserved the viable bacteria and the microbial thermolabile metabolites.

### Feeding and recording of piglet performance

This study was a feeding trial carried out on a commercial pig farm (Kawiks Farm, Patoki 23. 98– 170 Widawa. Province. Lodz city, Poland) in 2018, where groups of piglets were weaned one day a week over a 5-week period. The trial procedure and sample collection were approved by the Local Ethical Commission of Olsztyn University of Life Sciences (Olsztyn, Poland) with respect to experimentation and care of animals under study. A total of 690 piglets were tested under three different feeding regimens (230 piglets per feeding treatment) from 28 days of age (10 days before weaning) until 85 days of age when piglets exited the nursing unit. One group was a control group fed a basal feed according to Danish nutritional recommendations (Tybirk 2015), and the other two groups received supplementation of 2.5% or 5% FRS to the basal feed (feed dry matter basis) (Table 1). Piglets on each dietary regime were housed in nursing pens holding an average of 48 animals per pen. Each dietary treatment was repeated 5 times (1 repetition per experimental week and 1 pen representing a repetition) and the control was repeated 4 times. None of the diets included growth promoters, prescription antibiotics or zinc oxide. Piglets that experienced diarrhea or any other serious health conditions were removed from the experiment and treated elsewhere and counted as piglets that did not complete the experiment. Feed and fresh water were supplied *ad libitum* throughout the experiment.

**Table 1.**
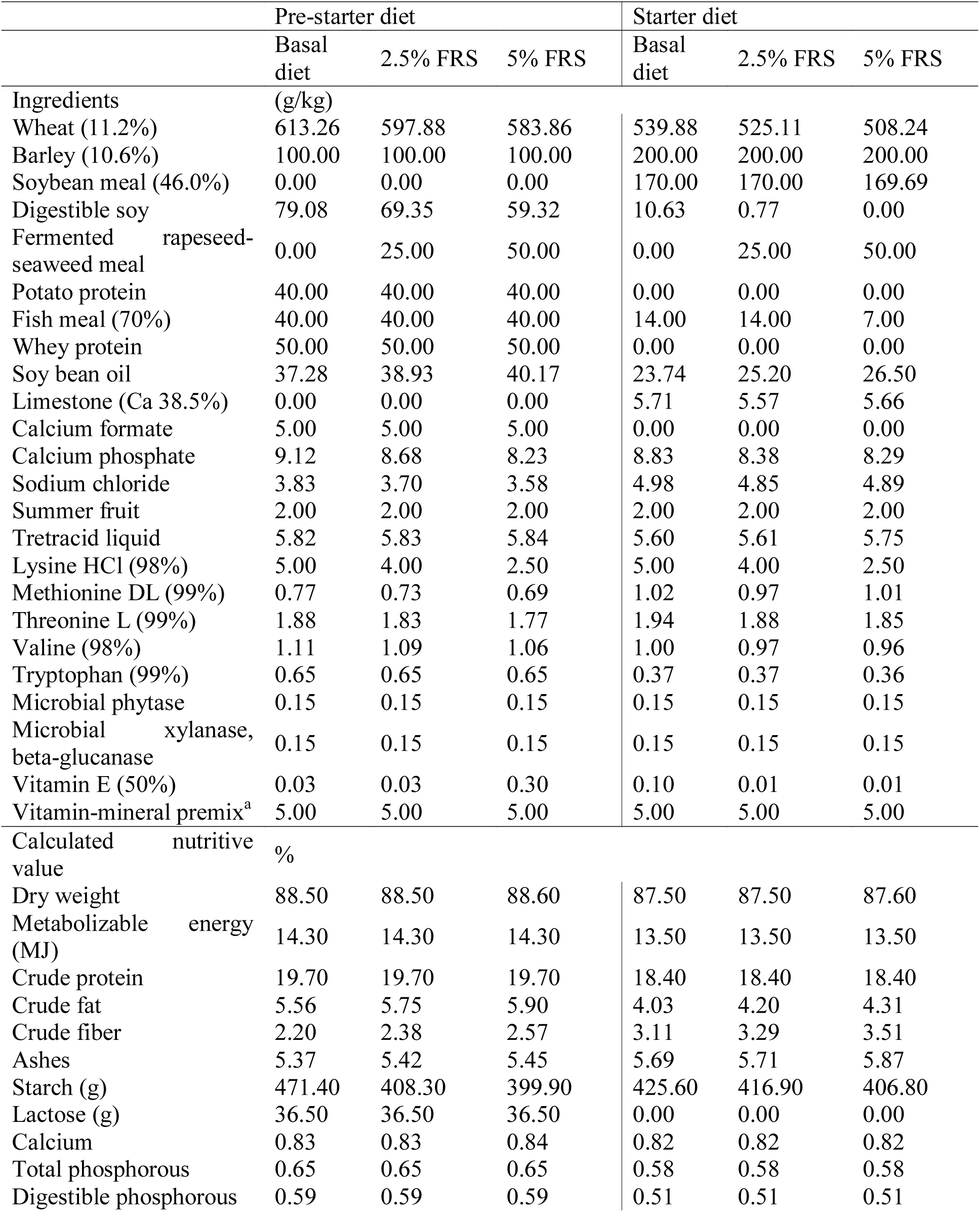
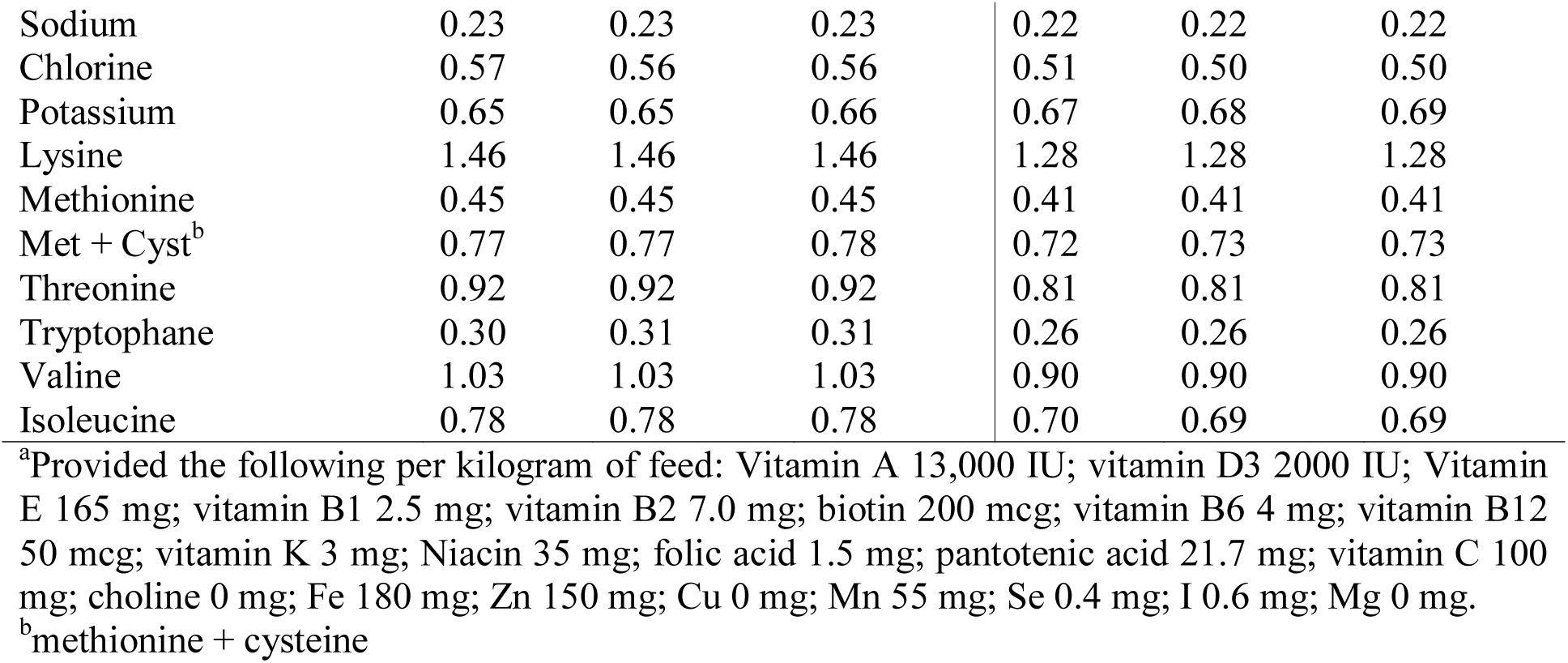
Feed formulations used for the experiment.

Litter weights during the nursing period were recorded every week, and feed intake was recorded daily. Performance indicators such as body weight, ADFI (average daily feed intake), average daily weight gain (ADG), feed conversion ratio (FCR) and completion rate were calculated at the pen level as previously outlined (Satessa et al. 2020a).

### Biological sample collection

A total of 10 piglets from each treatment (5 in each of two experimental weeks) were randomly selected and euthanized 3 weeks after weaning. The animals were euthanized by stun gunning with a captive bolt immediately followed by de-bleeding at the farm slaughtering facilities under strict sanitary regulations. Whole blood samples and serum for clinical analysis, the digesta from the colon for microbiome analysis, and jejunum and colon tissues for histopathological analyses, were collected in that order immediately after slaughtering.

A blood sample from each piglet was deposited in a tube with the anti-coagulant EDTA and preserved on ice until taken to the laboratory, where it was stored at 2-8 °C until analysis. Another blood sample was collected in a tube without anticoagulant, and serum separated by centrifugation, which was then stored at −20 °C until analysis. Gut tissues and colon contents were sampled after opening of the abdominal wall, and the stomach, small and large intestines were occluded at both ends and removed. Approximately 2 cm^3^ of colon content was collected from the apex of the ascending spiral of the colon with a sterile spatula and deposited in cryotubes with RNAlater™ (Sigma-Aldrich, Munich, Germany). Tubes with colon contents were kept at room temperature for less than 24 h, followed by cryopreservation in the laboratory. Tissue samples (approximate 2 cm long) of the whole transection of the jejunum and colon were excised and carefully rinsed from gut contents by flushing with saline (0.9 % NaCl). For each tissue a sterilized blade was used. Tissues were preserved in 10% formaldehyde and kept at room temperature for no longer than 24 hours until further processing (Satessa et al. 2020b).

### Blood hematology, blood biochemistry and serum immunoglobulin analysis

Full blood counts (erythrocyte, hemoglobin, hematocrit, mean corpuscular volume, mean corpuscular hemoglobin, mean corpuscular hemoglobin concentration, red cell distribution width) and differential white blood cells count (platelets, leucocytes, lymphocytes, monocytes, neutrophils, eosinophils, basophils) were performed using a Sysmex XT 2000i analyzer (Sysmex Corporation, Kobe, Japan). Serum analysis measured concentrations of the following, using standardized quantification methods: alanine aminotransferase, glutamic pyruvic transaminase, aspartate aminotransferase, glutamic-oxaloacetic transaminase, lactate dehydrogenase, lysozyme, glucose, total protein, blood urea nitrogen, uric acid, phosphorous, total cholesterol, triglycerides, low density lipoprotein, high density lipoprotein and immunoglobulin G (IgG) according to previously described procedures (Satessa et al. 2020a).

### Histological morphometric analysis of intestinal tissues

The histological analysis of mid-jejunal and colonic tissues was conducted by a commercial analytical laboratory (ALAB Weterynaria, Warsaw, Poland) according to previously described procedures (Satessa et al. 2020a). In short, tissue sections fixed in 10% formaldehyde were dehydrated by means of graded ethanol and xylene baths and embedded in paraffin wax, and 3-4 µm section were then stained with haematoxylin and eosin. Histopathological evaluations (at different lens magnifications) measured gut-associated lymphoid tissue (GALT), intraepithelial lymphocytes (IELs) and lymphatic infiltration of the stromal mucosa (stromal lymphocytes, SL) counts. In the GALT scoring, the numbers of lymphoid follicles per millimeter square were counted. For IEL scoring, the following scale was used: 0-normal (0-10 IELs/100 enterocytes), 1-low (10-15 IELs/100 enterocytes), 2-moderate (15-20 IELs/100 enterocytes; this level suggests chronic subclinical inflammation, where the intestinal-blood barrier may be damaged; weak lymphocytic inflammation), 3-severe (> 20 IELs/100 enterocytes; this level indicates chronic inflammation with infiltration damaging the epithelium and intestinal-blood barrier; moderate lymphocytic inflammation). For SL, the visual scoring scale was: 0-normal (single lymphocytes in stromal connective tissues of villus and crypts), 1-low (increased number of lymphocytes, but no damage to the stroma structures), 2-moderate (abundant infiltration of lymphocytes in stroma, damaging blood vessel walls, connective tissue fiber, reducing visibility of stroma structures), 3-severe (lymphocyte infiltration completely disrupts and conceals the stroma). In a blinded fashion, 10 fields of view per piglet at 4× magnification were used for evaluation of GALT structures and numbers of lymphoid follicles. IEL and SL were evaluated at 40× magnification. The analysis used a standard light microscope Olympus BX41 and Cell Sens software (Olympus Corporation, Tokyo, Japan). The gut tissues samples which could not reach the requirements for histological analysis were discarded, which resulted in 9, 8, 10 piglets included in analysis for the basal diet, 2.5% and 5% FRS groups, respectively.

### 16S rRNA gene amplicon sequencing of colon content

Collected colon contents was stored at −60 °C prior to the analysis. Two types of 16S rRNA gene amplicon sequencing strategies were adopted to characterize the prokaryotic community: Illumina, NextSeq (Illumina, CA, USA) and MinION (Oxford Nanopore Technologies, Oxford, UK). The genomic DNA was extracted using Bead-Beat Micro AX Gravity Kit (A&A Biotechnology, Gdynia, Poland) according to the manufacturer’s instruction. DNA concentration and purity were measured using NanoDrop ND-1000 spectrophotometer (Saveen and Werner AB, Sweden).

Extracted DNA was diluted to 10 ng/µL prior to library preparation. The V3 hypervariable region of 16S rRNA gene was amplified and sequenced with Illumina technology as previously described (Krych et al. 2018). Near full-length 16S rRNA gene amplicons were amplified and sequenced with ONT targeting V1-V8 hypervariable region using following primers: ONT_27Fa: GTCTCGTGGG CTCGGAGATG TGTATATAGA TCGCAGAGTT TGATYMTGGCTCAG; ONT_27Fb: GTCTCGTGGG CTCGGAGATG TGTATATAGA TCGCAGAGTT TGATCCTGGCTTAG and ONT_1540_R: GTCTCGTGGG CTCGGAGATG TGTATACTCT CTATTACGGY TACCTTGTTACGACT. Custom designed barcoding system was developed to tag encode up to 96 samples during the second round of PCR, and the PCR primer sequence is given in Appendix Table 1. The PCR1 reaction mix contained 5µl of PCRBIO buffer and 0.25 µL PCRBIO HiFi polymerase (PCR Biosystems Ltd, London, United Kingdom), 1 µL of primers mix (5 µM of ONT_27Fa and ONT_27Fb, and 10 µM of ONT_1540_R, see above), 5 µL of genomic DNA (∼10ng/µL) and nuclease-free water to a total value of 25 µL. The PCR thermal conditions were as follows: denaturation at 95°C for 5 min; 33 cycles of 95°C for 20 s, 55°C for 20 s and 72°C for 45 s; followed by final elongation at 72°C for 4 min.

PCR products were verified by agarose gel electrophoresis and then subjected for barcoding (PCR2). The PCR2 mix composed of 5 µL PCRBIO buffer, 0.25 µL PCRBIO HiFi polymerase (PCR Biosystems Ltd, London, United Kingdom), 2 µL of barcode primers (5 µM) (Appendix Table 1) 1 µL of PCR1 template and DEPC water up to 25 µL. The PCR2 thermal conditions were as follows: denaturation at 95 °C for 2 mins; 13 cycles of 95 °C for 20 s, 55 °C for 20 s, 72 °C for 40 s; final elongation at 72 °C for 4 mins. The final PCR products were purified using AMPure XP beads (Beckman Coulter Genomic, CA, USA) and pooled in equimolar concentrations. The pooled barcoded amplicons were subjected to 1D genomic DNA by ligation protocol (SQK-LSK109) to complete library preparation for MinION sequencing. Approximate 0.2 μg of amplicons were used for the initial step of end-prep. And 40 ng of prepared amplicon library was loaded on a R9.4.1 flow cell.

### Sequencing data analysis

The raw Illumina derived dataset containing pair-ended reads with corresponding quality scores were merged and trimmed using fastq_mergepairs and fastq_filter scripts implemented in the USEARCH pipeline as described previously (Krych et al. 2018). Purging the dataset from chimeric reads and constructing zero radius Operational Taxonomic Units (zOTU) was conducted using the UNOISE. The Greengenes (13.8) 16S rRNA gene collection was used as a reference database (DeSantis et al. 2006).

Data generated by MinION were collected using MinKnow software v19.06.8 (https://nanoporetech.com). The Guppy v3.2.2 basecalling toolkit was used to base call raw fast5 to fastq (https://nanoporetech.com). Porechop v0.2.2 was used for adapter trimming and sample demultiplexing (https://github.com/rrwick/Porechop). The Porechop adapter list was (adapters.py) edited accordingly and is given in Appendix Table 1. Sequences containing quality scores (fastq files) were quality corrected using NanoFilt (q ≥ 10; read length > 1Kb). Taxonomy assignment of quality corrected reads against Greengenes (13.8) database was conducted using uclast method implemented in parallel_assign_taxonomy_uclust.py QIIME (v1.9.1). The uclust settings were tuned on mock communities (--similarity 0.8; min_consensus_fraction 0.51) assuring annotations to the lowest taxonomic level with no false positive annotations. The settings allowed it to treat individual Oxford Nanopore Technologies-Amplicon Sequence Variant (ONT-ASV) as individual “seeds”. Reads classified to at least phylum level were subjected for further analysis.

### Statistics

All the statistical analysis concerning phenotype data was performed with R (v3.6.2). The difference of piglets performance on the pen level was evaluated using linear mixed model as previously outlined (Satessa et al. 2020a) and orthogonal polynomial contrast was used to appreciate the effect on increasing dose of the FRS (0%, 2.5%, 5%). For histology and hematology data, Wilcoxon rank test was used to evaluate statistical significance between different treatment groups.

For microbiome analysis, QIIME2 (Bolyen et al. 2019) (v2018.11) combined with R packages (ggplot2, vegan, corrplot, Rhea, rstatix, vennDiagram) were used. Three samples were removed due to inadequate library size (< 1000 counts), which resulted in 9, 8, 10 piglets included for basal diet, 2.5% and 5% FRS group, respectively. Considering both types of amplicon profiling, all the samples were summarized at the L7 levels (species) and rarefied to the same sequencing depth (11000 reads/sample) for alpha and beta diversity calculations on both platforms. The standard rarefaction at ASV level was conducted on Illumina data as comparison for rarefaction on species level. Principal coordinate analysis (PCoA) plots were conducted on binary Jaccard and Bray Curtis distance metrics, and PERMANOVA was performed to check differences between groups and *p* values were FDR adjusted after pairwise comparison. ANCOM (Mandal et al. 2015) was adopted to detect statistical differences among groups at summarized L7 level using default settings in QIIME2. For all taxa found to differ between treatments by ANCOM, Wilcoxon rank-sum test was conducted for pairwise comparison using the respective relative abundances.

Phenotypic (including clinical and immunological) data were integrated with microbiota data by Pearson’s correlation analysis using Rhea (Lagkouvardos et al. 2017). Rare microbial appearances at species level were removed with a cutoff of mean relative abundance > 0.1% and minimal presence among 30% of samples. Zeros were regarded as NA. Centered log-ratio transformation was conducted in both the microbial relative abundance and phenotypic data. The correlation matrix was visualized with R package corrplot using the same defined threshold (*p* < 0.05) for both platforms.

## Results

### Piglet performance, blood hematology, blood biochemistry and systemic immunoglobulin

Piglets fed with 2.5% FRS had numerically, but not significantly increased body weight by the end of the experiment (85 days of age) in comparison with those fed the basal diet. Further, increasing the dose of FRS to 5% did not positively influence the final body weight of animals (Table 2). No significant differences in average daily weight gain (ADWG), average weight gain (AWG), or feed conversion ratio (FCR) were recorded between the three feeding regimens. The completion rate for piglets in the experiment (ie. not dead or removed due to need for antibiotics treatment) did not differ between treatment groups. In the sub-group of piglets euthanized 3 weeks after weaning, we found no statistical differences in blood hematology, blood chemistry and systemic immunoglobulin parameters among treatment groups, except for the blood urea nitrogen (BUN) and mean corpuscular characteristics (Appendix Fig. 1). Compared with piglets fed the basal diet, 2.5% FRM reduced plasma level of BUN, but increased mean corpuscular volume and mean corpuscular hemoglobulin in blood. Piglets fed with 5% FRM had a similar tendency, but only BUN level was significantly lowered relative to the basal diet.

**Table 2.** Performance of piglets subjected to three dietary regimens.

| Parameters | Basal diet<br>(0% added) | 2.5% FRS<br>(2.5% added) | 5% FRS<br>(5% added) | SEM | P-value |  |  |
| --- | --- | --- | --- | --- | --- | --- | --- |
|  |  |  |  |  | TG | TG |  |
|  |  |  |  |  |  | L | Q |
| Weaning weight<br>(kg, at d 28) | 6.07 ± 0.33 | 6.24 ± 0.34 | 5.99 ± 0.66 |  |  |  |  |
| Body weight, at age of interest (kg) |  |  |  |  |  |  |  |
| 42 | 6.77 | 7.00 | 6.55 | 0.406 | 0.579 | 0.890 | 0.428 |
| 49 | 8.24 | 8.75 | 8.41 | 0.418 | 0.563 | 0.746 | 0.314 |
| 77 | 20.5 | 21.6 | 20.2 | 1.18 | 0.537 | 0.842 | 0.341 |
| 85 | 23.2 | 25.1 | 23.7 | 1.82 | 0.467 | 0.823 | 0.269 |
| ADFI (kg/day) |  |  |  |  |  |  |  |
| 28 – 42 days | 0.162 | 0.183 | 0.166 | 0.015 | 0.437 | 0.830 | 0.231 |
| 28 – 49 days | 0.233 | 0.232 | 0.230 | 0.020 | 0.985 | 0.900 | 0.967 |
| 28 – 85 days | 0.516 | 0.572 | 0.546 | 0.037 | 0.418 | 0.477 | 0.236 |
| 50 – 77 days | 0.626 | 0.661 | 0.632 | 0.031 | 0.531 | 0.879 | 0.299 |
| 50 – 85 days | 0.671 | 0.763 | 0.719 | 0.058 | 0.365 | 0.460 | 0.199 |
| ADG (kg/day) |  |  |  |  |  |  |  |
| 28 – 42 days | 0.049 | 0.063 | 0.049 | 0.025 | 0.835 | 0.993 | 0.574 |
| 28 – 49 days | 0.103 | 0.122 | 0.118 | 0.017 | 0.681 | 0.506 | 0.499 |
| 28 – 85 days | 0.306 | 0.339 | 0.316 | 0.034 | 0.584 | 0.790 | 0.343 |
| 50 – 77 days | 0.439 | 0.456 | 0.420 | 0.037 | 0.623 | 0.702 | 0.476 |
| 50 – 85 days | 0.423 | 0.467 | 0.431 | 0.045 | 0.486 | 0.993 | 0.574 |
| FCR |  |  |  |  |  |  |  |
| 28 – 42 days | 6.78 | 7.42 | 7.13 | 3.12 | 0.938 | 0.878 | 0.740 |
| 28 – 49 days | 2.37 | 2.12 | 2.09 | 0.311 | 0.681 | 0.486 | 0.683 |
| 28 – 85 days | 1.80 | 1.68 | 1.76 | 0.147 | 0.584 | 0.834 | 0.515 |
| 50 – 77 days | 1.44 | 1.45 | 1.55 | 0.084 | 0.485 | 0.365 | 0.597 |
| 50 – 85 days | 6.78 | 7.42 | 7.13 | 3.12 | 0.938 | 0.878 | 0.740 |
| Completion rate (%) |  |  |  |  |  |  |  |
| 28 – 42 days | 95.2 | 96.9 | 96.1 | 2.10 | 0.773 | 0.74 | 0.504 |
| 28 – 49 days | 94.1 | 95.1 | 94.3 | 2.29 | 0.847 | 0.921 | 0.601 |
| 28 – 85 days | 94.8 | 94.4 | 89.1 | 3.60 | 0.472 | 0.328 | 0.564 |

ADFI = average daily feed intake; ADG = average daily gain; FCR = feed conversion ratio; TG = treatment group; L = linear effect; Q = quadratic effect; The significance between different dosage group is based on the result of orthogonal contrasts.

**Fig. 1.**
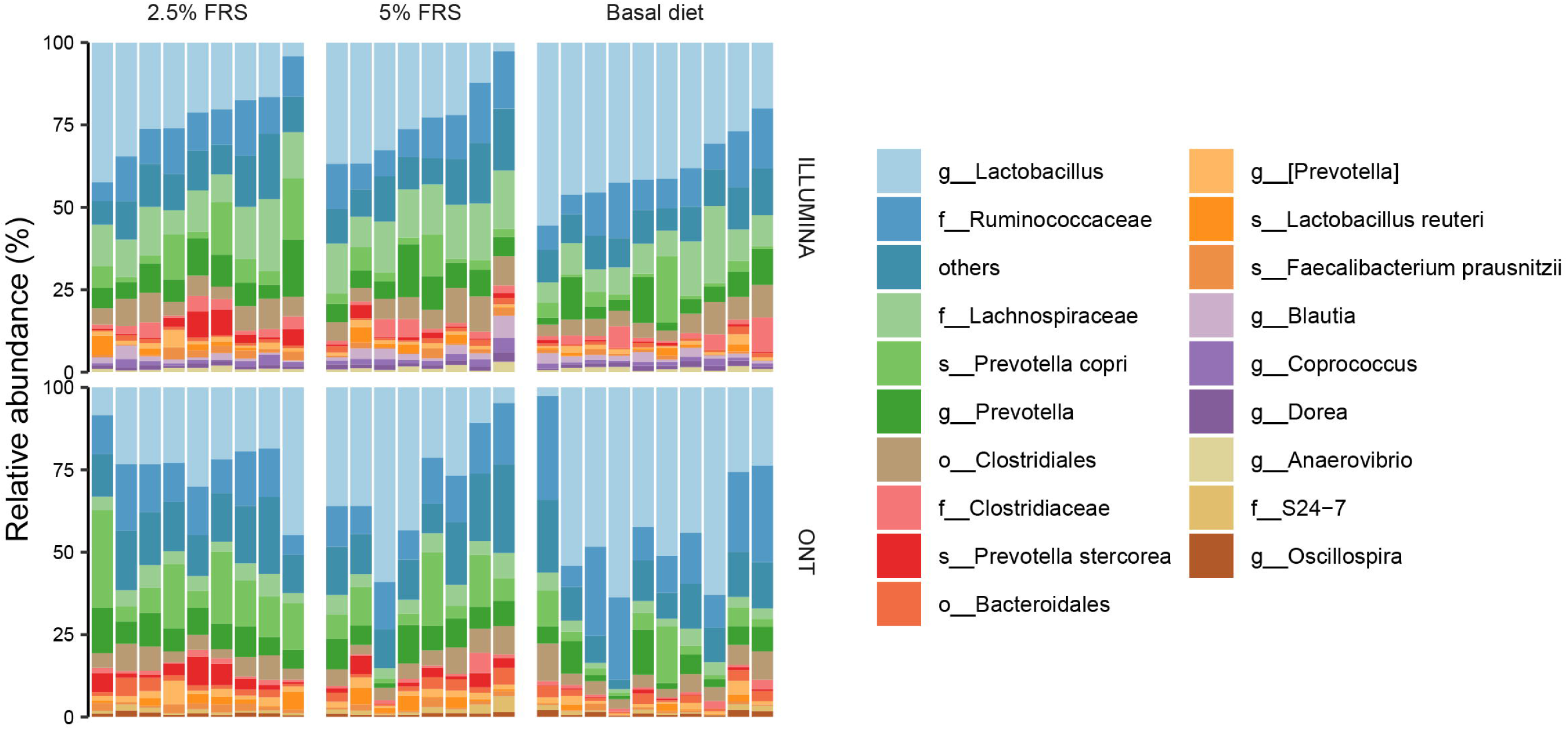
Relative abundance of prokaryotes based on 16S rRNA gene amplicon sequencing using either the Illumina or the Oxford Nanopore Technologies platform. Respectively n= 9, 8, 10 for basal diet with 0%, 2.5% and 5% added FRS (fermented rapeseed-*Sacharina latissima*-*Ascophillum nodossum*).

### High level of accordance between the short and long amplicon sequencing strategies

Two different sequencing strategies were applied. Illumina NextSeq-based amplicon sequencing of the 16S rRNA gene V3 variable region (Illumina V3) and ONT based sequencing of V1-V8 variable regions (ONT V1-V8). Out of 99 unique taxonomic groups discovered with both methods, 78 were shared between the two sequencing methods (Appendix Fig. 2A). The accordance between the two sequencing strategies was further improved after abundance threshold adjustment (Appendix Fig. 2B-D). The level of similarity between taxa identified with both sequencing strategies reached 100% when analyzing taxonomic groups with relative abundance above 3% (Appendix Fig. 2E) and overall with very good correlation between the two methods (Appendix Fig. 2F). Both methods revealed that the most dominant bacterial groups belonged to genus *Lactobacillus* and families: *Ruminococcae* and *Lachnospiraceae* independent of treatment (Fig 1).

**Fig. 2.**
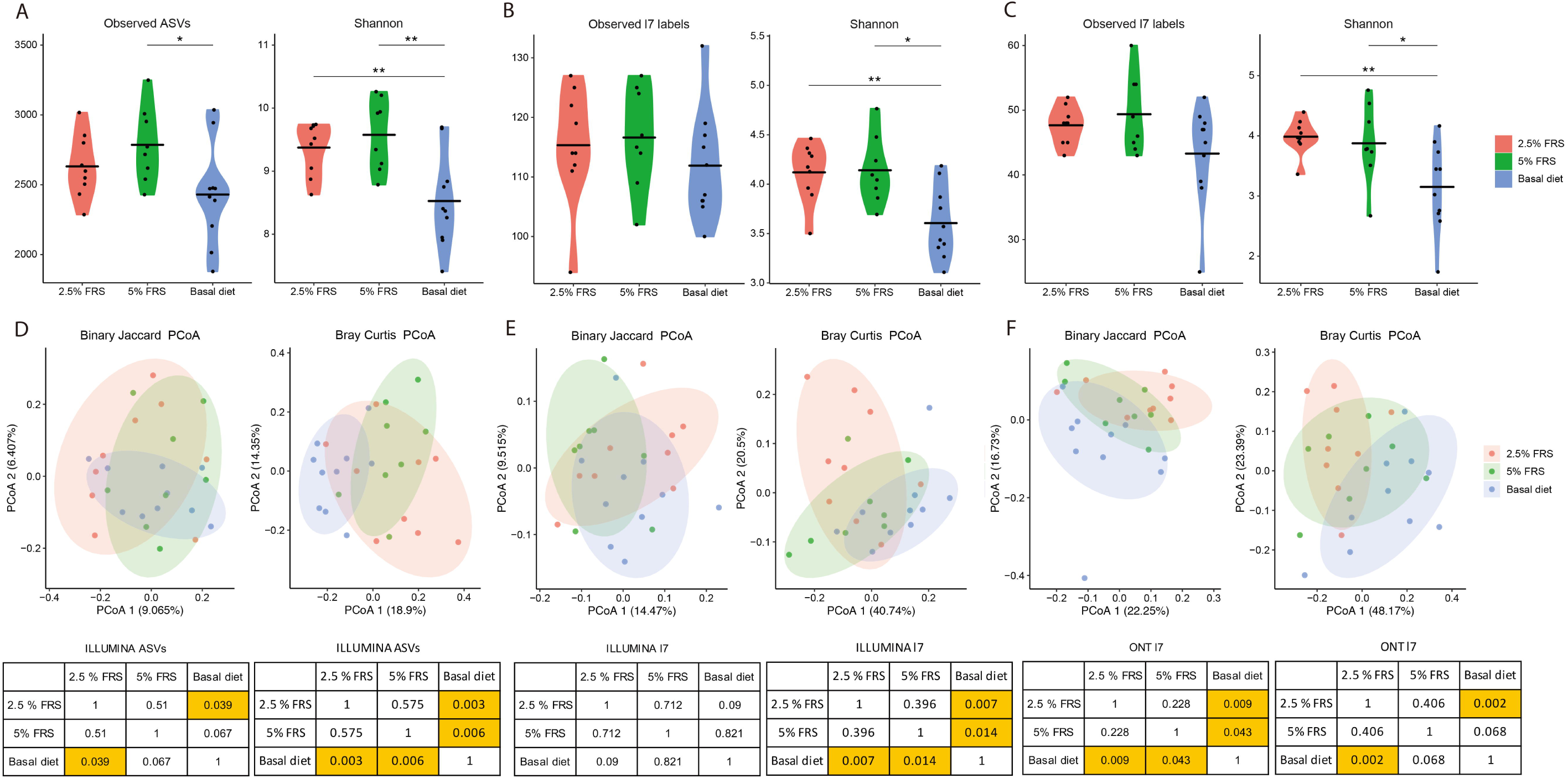
Alpha and beta diversity analysis of 16S rRNA gene amplicon sequencing data. Observed ASVs and Shannon index based on rarefied ASV table with Illumina sequencing on V3 region (Illumina V3) (A); Observed features and Shannon index based on species-level summarized table with Illumina sequencing on V3 region (B) and Oxford Nanopore sequencing on V1-V8 region (ONT V1-V8) (C). The mean value for each group is marked as a bold line respectively. Respective PCoA plot of binary Jaccard and Bray Curtis distance metrics based on rarefied ASV table (Illumina V3) (D), species-level summarized table (Illumina V3) (E) and species-level summarized table (ONT V1-V8) (F). The ellipses show respective 80% confidential area following multivariate t-distribution. Respectively n= 9, 8, 10 for basal diet with 0%, 2.5% and 5% added FRS (fermented rapeseed- *Sacharina latissima*- *Ascophillum nodossum*). For pairwise Wilcoxon test on alpha diversity, the labels of *, ** represent *p* < 0.05, < 0.01 respectively. For PERMANOVA tests, *p* values below 0.05 were heighted in yellow.

### Supplementation with FRS induced clear changes in the gut microbiota composition

Data generated with both sequencing approaches revealed differences in gut microbial diversity in piglets 3 weeks after weaning according to dietary treatment. FRS addition in the feed resulted in increased Shannon index but not Observed feature index values. The effect was consistent, when the analysis was performed based on the non-summarized ASV-table (Fig 2A) and summarized to the species level table (Fig. 2B) from Illumina data, and when summarized to the species level from ONT data (Fig. 2C). Increasing the dietary inclusion of FRS from 2.5% to 5% of DM did not cause significant changes in the alpha diversity indices between the two groups (Fig. 2A-C). Beta diversity analysis based on binary Jaccard distance (qualitative) and Bray Curtis dissimilarity (quantitative) showed that introduction of FRS in the feed influenced colon microbiota composition. Surprisingly, the changes were more pronounced within the 2.5% FRS supplemented group compared to the 5% FRS (Fig. 2D-F).

### Prevotella stercorea and Faecalibacterium prausnitzii are positively correlated with colon mucosa thickness but negatively correlated with systemic IgG levels

The relative abundance of *Prevotella stercorea* and *Mitsuokella spp*. were increased in the 2.5% FRS-feeding group compared to basal diet and for most comparisons also in the 5% FRS group as well (Fig. 3A-D). The relative abundance of *Prevotella stercorea* (Illumina V3) was shown to be positively correlated with the colon mucosa thickness and negatively correlated to the blood leucocytes counts and serum IgG level (Fig. 4A), while this observation was only near-significant using ONT (Fig. 4B).

**Fig. 3.**
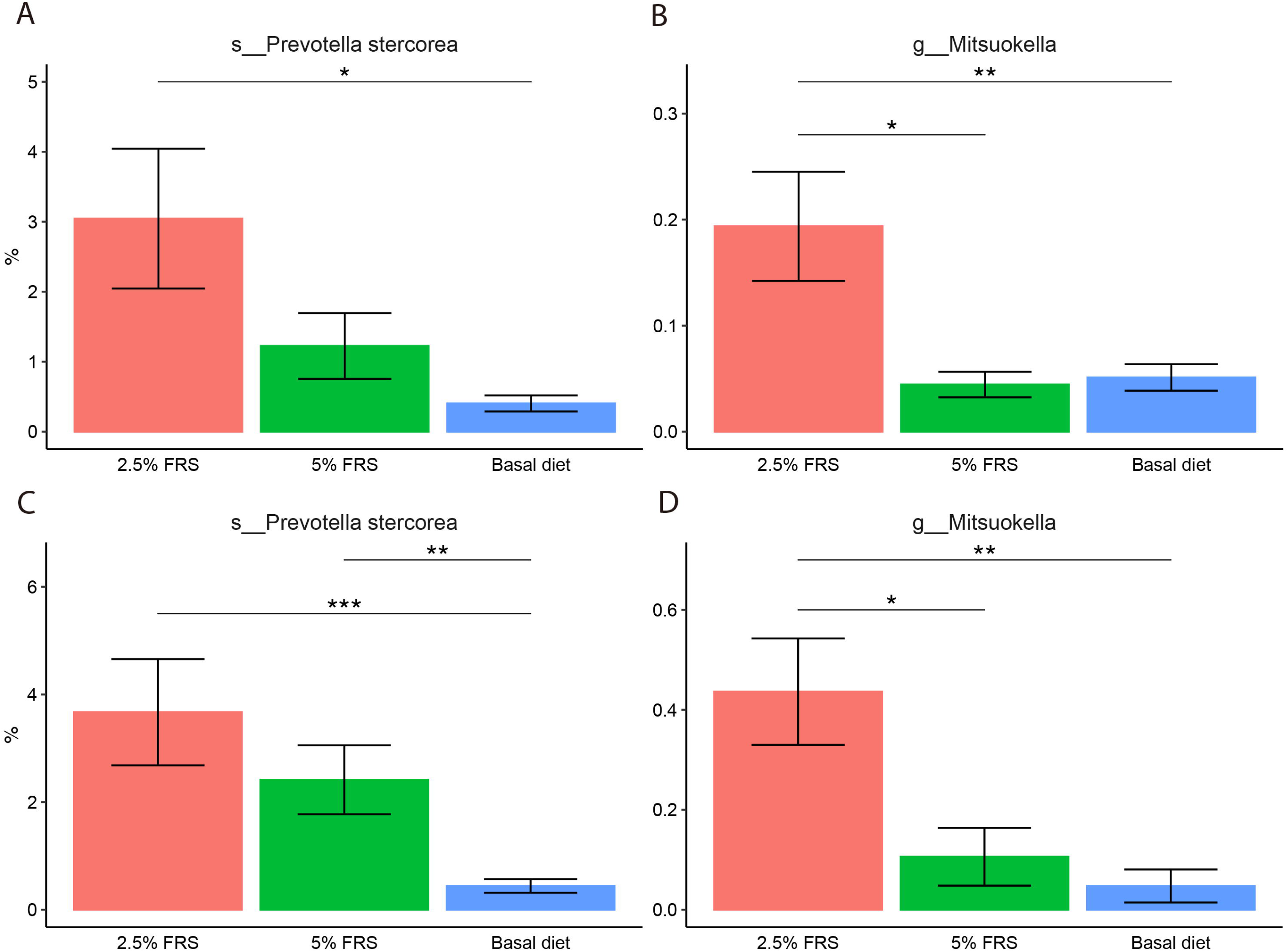
*Prevotella stercorea* and *Mitsuokella* spp. showed different colonized abundance in gut between feeding regime groups by both Illumina sequencing on V3 region of 16S rRNA gene (A, B) and Oxford Nanopore sequencing on V1-V8 region of 16S rRNA gene (C, D). Data in the bar plot was presented as mean value and SEM error bar. Respectively n= 9, 8, 10 for basal diet with 0%, 2.5% and 5% added FRS (fermented rapeseed- *Sacharina latissima*- *Ascophillum nodossum*). The labels of *, **, *** represent *p* < 0.05, < 0.01, < 0.005 respectively.

**Fig. 4.**
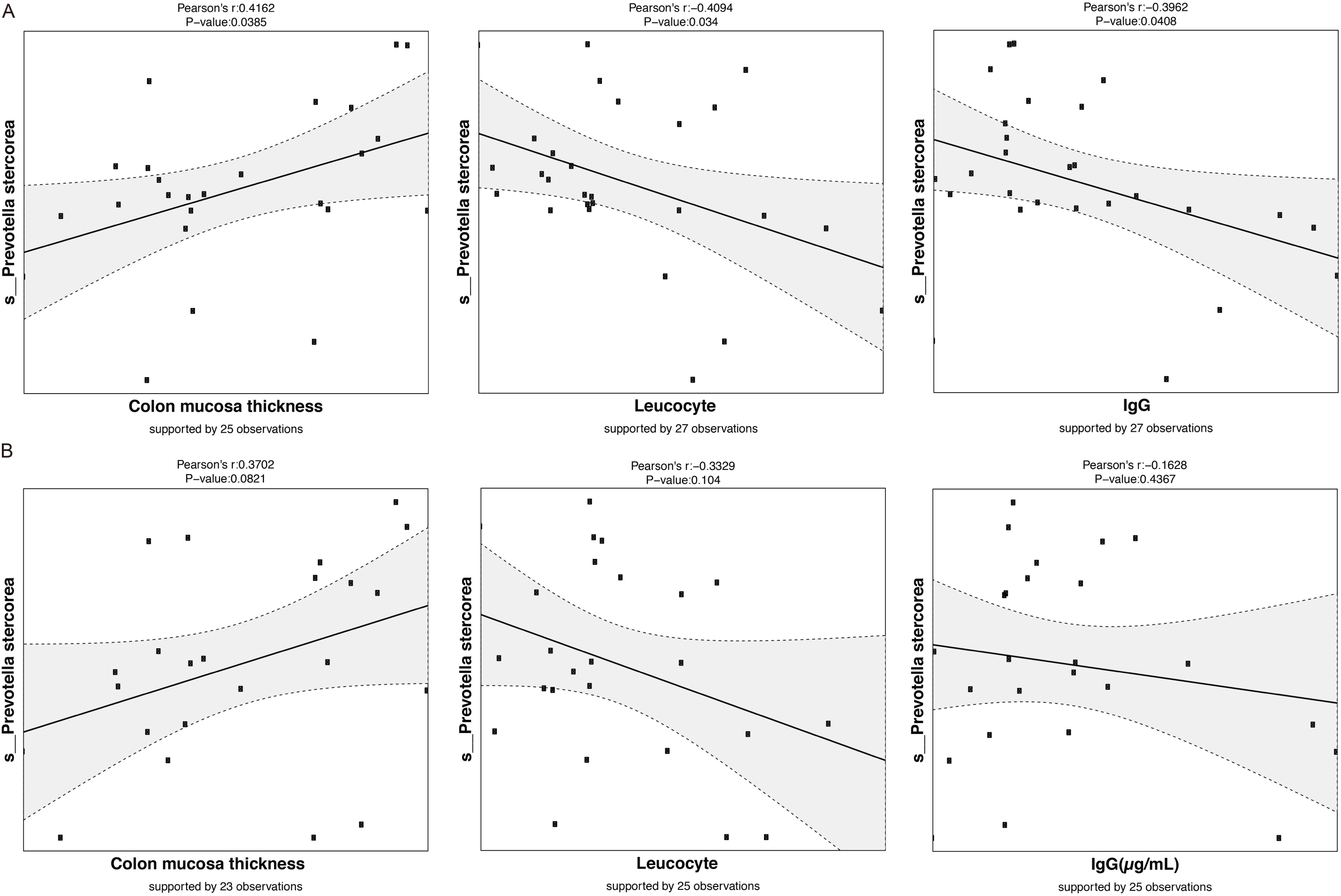
*Prevotella stercorea* significantly correlated with colon mucosa thickness, serum leucocyte and IgG with data from V3 region 16S rRNA gene sequencing (A), similar trend yet not significant was observed using V1-V8 region 16S rRNA gene sequencing (B). Data was centered log-ratio transformed for Pearson’s correlation.

There were 83 significant correlations between bacteria relative abundance verified with Illumina V3 region and indicators of systemic or intestinal immunomodulation or intestinal histopathological parameters, while ONT resulted in 114 significant correlations. Although many matching results (associations and trends) could be found comparing correlation results from both methods, only one taxon, *Faecalibacterium prausnitzii* presented 100% accordance in correlations from the two sequencing strategies. *F. prausnitzii* relative abundance correlated positively with colon mucosa thickness and negatively with serum levels of aspartate aminotransferase, lactate dehydrogenase and IgG (Appendix Fig. 3).

### Supplementation with FRS protected the gut health and alleviated inflammation

The morphological characteristics of intestinal tissues obtained from piglets in the different treatment groups are shown in Fig. 5A. All animals fed 2.5%, 5% FRS or the basal diet presented normal ranges for heights and structures of villi and intestinal crypts. The continuity and height of the jejunal and colonic epithelium was more pronounced in both FRS groups compared to the piglets on the basal feed. Histological evaluation of tissues was performed for lymphocytic infiltration, GALT structures and colon mucosa thickness. In the jejunal epithelium and stroma, the piglets fed with FRS had reduced intraepithelial lymphocytes (IEL) and stromal lymphocytes (SL) infiltration compared to piglets fed the basal diet, with 2.5% FRS showing the best effect (Fig. 5B). We observed similar tendency of alleviated focal inflammation in the colon tissues of 2.5% FRS, but no significant difference was found between treatment groups (Fig. 5C). Diffuse lymphoid follicles at the base of the mucosa were visible with normal size and structure in jejunum and colon. No clear stimulation of lymphoid follicles was observed in any of the three dietary groups. Neither did FRS supplementation result in differences in the number of jejunal and colonic GALT structures relative to the basal diet (Fig. 5D). The histological evaluation did not indicate damage of the intestinal epithelial barrier in any group but the mucous membrane was higher with significantly deeper intestinal crypts within the 2.5% FRS group compared to the basal diet group (Fig. 5E).

**Fig. 5.**
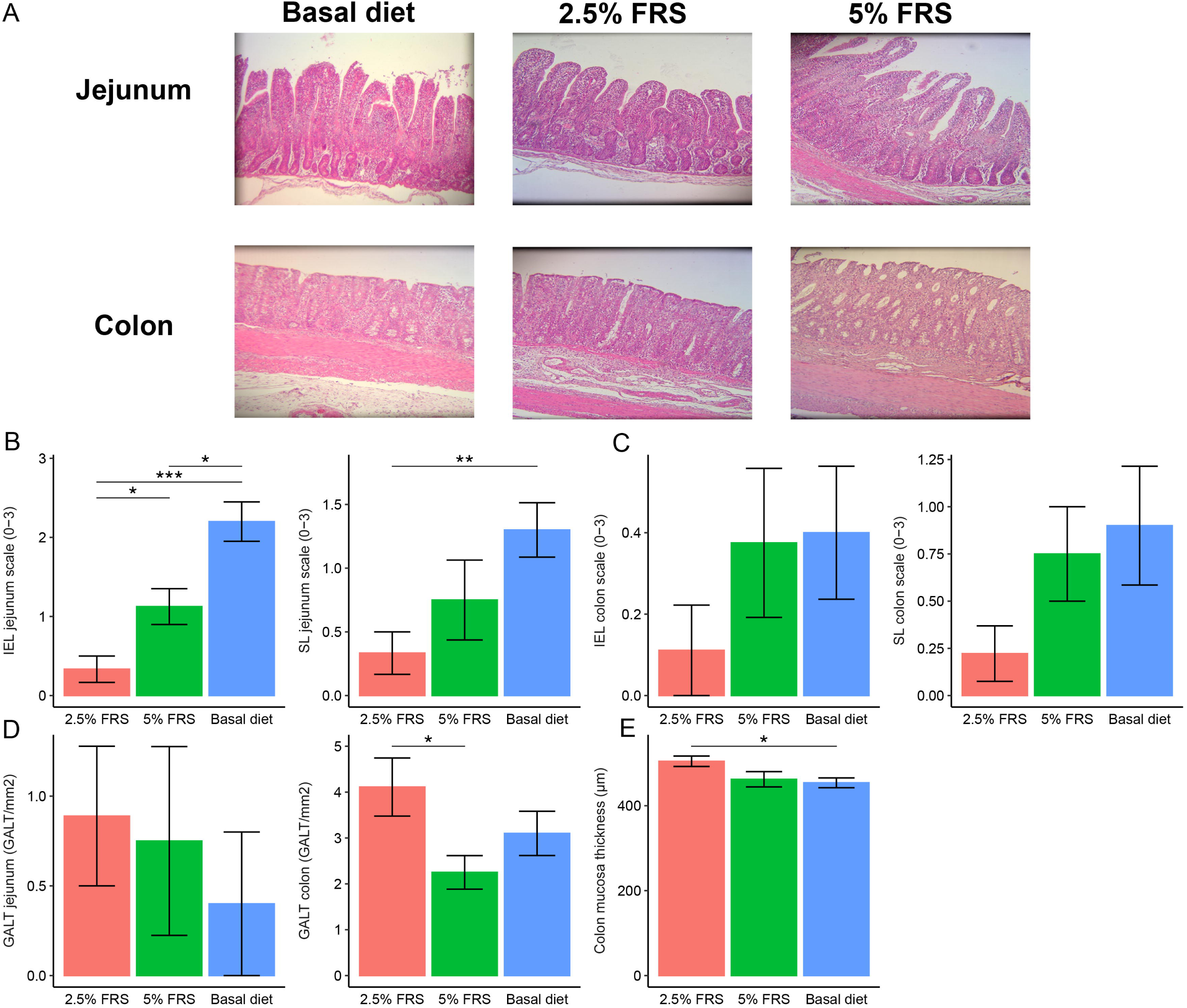
2.5% FRS feeding protected the gut health of piglets with alleviated inflammation symptoms and enhanced mucosa barrier in gut. Histopathological micrograph of jejunum and colon under 10x magnification (A), levels of inflammation measured by the number of lymphocytes infiltrated at the epithelium (IEL) and at the stroma (SL) of jejunum (B) and colon (C), levels of immune capacity estimated by numbers of lymphoid follicles per squared millimeter in jejunum and colon (D) and mucosa thickness in colon (E). Respectively n= 9, 8, 10 for basal diet with 0%, 2.5% and 5% added FRS (fermented rapeseed- *Sacharina latissima*- *Ascophillum nodossum*). The labels of *, **, *** represent *p* < 0.05, < 0.01, < 0.005 respectively.

## Discussion

In modern pig production, the weaning of piglets is usually conducted at an early age causing significant stress due to changes in diet, environment and social groups. Hence, many weaned piglets experience intestinal and immune dysfunction, elevated risk of infection with enteric pathogens and hence diarrhea, and lowered weight gain due to reduced feed intake and poorer utilization of ingested nutrients (Lallès et al. 2007). Reduced weight gain and higher mortality rate among weaned piglets are highly undesirable from a production efficiency point of view. Preventive measures, such as use of in-feed antibiotics have been banned by the European Union (EU) back in 2006, while the commonly used zinc oxide (ZnO) will be banned in 2022. Therefore, there is an urgent need for development of alternative preventive strategies in order to sustain performance and a healthy gut of weaned piglets. We have previously reported that FRS fed to weaned piglets improved jejunal villus development, stimulated colon mucosal development and reduced signs of intestinal inflammation (Satessa et al. 2020a). Since FRS was demonstrated to be effective without in-feed ZnO, we further investigated in the present study the dose-dependent influence of FRS on gut microbial composition and its plausible link with indicators of intestinal and systemic immune function.

To study the gut microbial composition, we have used and compared the performance of two sequencing strategies. The widely used short-read 16S rRNA gene amplicon sequencing performed on an Illumina-platform was compared with the less commonly used near-full length 16S rRNA gene sequencing method by ONT. The two strategies showed satisfying accordance and allowed to draw the same overall conclusions, including those identified at the species level, which is challenging even when different hypervariable regions of 16S rRNA gene within the same sequencing method are compared (Bukin et al. 2019) (Yang et al. 2016) (Kerrigan et al. 2019). Data generated with both sequencing strategies confirmed significant changes in gut microbiota qualitative and quantitative characteristics in response to dietary FRS supplementation. The effect was more pronounced for the 2.5% FRS group compared to the 5% FRS group. Piglets fed the 2.5% and 5% FRS supplemented diets had higher Shannon index indicating a more diverse and uniformly distributed gut microbiome than the piglets fed basal diet. The Observed features index was largely similar between all 3 groups. High microbial diversity is generally desirable, as it has been demonstrated to exclude pathogenic microbes, improve immune response and reduce necrotizing enterocolitis and post-weaning diarrhea incidences (Fouhse et al. 2016) (Khanna et al. 2016) (Dou et al. 2017). FRS supplementation led to increased relative abundance of *Prevotella stercorea* and *Mitsuokella* spp. relative to piglets fed the basal diet, an effect that was especially pronounced in the 2.5% FRS group. *Prevotella* is known to be the major contributor to the microbiome of post-weaned piglets due to ability to degrade plant fibers in the solid diet. The species *P. stercorea* has previously been described as a member of the healthy pig’s GM (Wang et al. 2019) as well as a potent producer of short-chain fatty acids (SCFA), when carbohydrates such as fibers are fermented in the hind-gut (De Filippo et al. 2010)’ (Chen et al. 2017b). Our data indicated that the abundance of *P. stercorea* correlated positively with colon mucosa thickness, which is not surprising, since *Prevotella* spp. are recognized colonizers of mucosal sites (Larsen 2017) and the produced SCFAs help maintain intestinal barrier function through providing energy resources and immunoregulatory regulation as well (Kiefer et al. 2006)’ (Chen et al. 2017a)’ (Spiljar et al. 2017). The negative correlation of *P. stercorea* with the serum levels of leucocytes and IgG could indicate that increased abundance of *P. stercorea* on the more fibrous FRS supplemented diet stimulated gut barrier and immune function, and consequently relieved the host inflammation. Previous studies also report that complex hemicelluloses and cellulose most likely enhances mucosal abundance of *P. stercorea* (Mann et al. 2014) (Mach et al. 2015).

*Faecalibacterium prausnitzii* is one of the main butyrate producers found in the gastro-intestinal tract (Barcenilla et al. 2000). Butyrate plays a vital role in gut physiology and gut health. It serves as a main energy source for the colonocytes and plays a protective role against inflammatory disease and colorectal cancer (Christl et al. 1996)’ (Archer et al. 1998). Many studies have linked reduced abundance of *F. prausnitzii* with different intestinal disorders, hence it has been proposed that *F. prausnitzii* may serve as a biomarker of gut health (Lopez-Siles et al. 2017). Our data showed that *F. prausnitzii* correlated positively with colon mucosa thickness and negatively with serum levels of two enzymes released from the liver e.g. aspartate aminotransferase (AST) and lactate dehydrogenase (LDH) as well as serum IgG. Increased serum level of the hepatic enzymes is a sign of liver malfunction while IgG is systemic indicator of inflammation status of the host. Hence, our findings are in line with studies demonstrating the ability of *F. prausnitzii* to reduce inflammation and improve the liver function in murine models (Munukka et al. 2017) (Fukui 2019) and humans.

Although the data on microbiota composition suggests there was no distinct impact with regards to the inclusion levels of FRS (2.5% versus 5%), it is important to note that significantly alleviated signs of lymphocyte invasion in jejunum and enhanced colon mucosa barrier function were observed solely in the group receiving 2.5% FRS. Moreover, the 5% FRS addition tended to numerically worsen production performance (ADG, FCR, completion rate). Possibly, if higher amounts of FRS are added, the piglets will also be exposed to more bioactive components from either the rapeseed or seaweed, which even though they are beneficial in lower amounts, could increase the physiological stress in the weaning period if added in relatively high amounts and be counterproductive to FCR in production. Young animals are usually more sensitive to these anti-nutritional factors, like glucosinolates (Tripathi and Mishra 2007) than older animals, and thereby our results suggest that the optimal inclusion level was reached at around 2.5% FRS.

## Conclusion

Our study demonstrates that dietary supplementation in postweaning piglets with 2.5% FRS led to significant improvements of gut health and induced favorable changes in gut microbiota composition, while further increasing the dietary addition to 5% FRS resulted in less pronounced effects. The 2.5% FRS supplementation induced significant changes in gut microbial composition expressed as increased microbial diversity index and elevated abundance of *Prevotella stercorea* and *Mitsuokella* spp. Although clear causality cannot be proven, we found clear correlations between the abundance of *Prevotella stercorea* and *Faecalibacterium prausnitzii* and biomarkers of reduced intestinal and systemic inflammation, improved liver function, and increased colon mucosal thickness.

## Acknowledgement

This work was supported by Bio-Based Industries Joint Undertaking under the European Union Horizon 2020 research and innovation program under grant agreement No 720755 (Macro Cascade project). Gizaw Dabessa Satessa held a PhD scholarship co-financed by a grant (file no. 5157-00003B) from the Innovation Fund Denmark and the University of Copenhagen, Denmark. Yan Hui was financed by the China Scholarship Council scholarship.

## Appendix

**Appendix Table 1.**
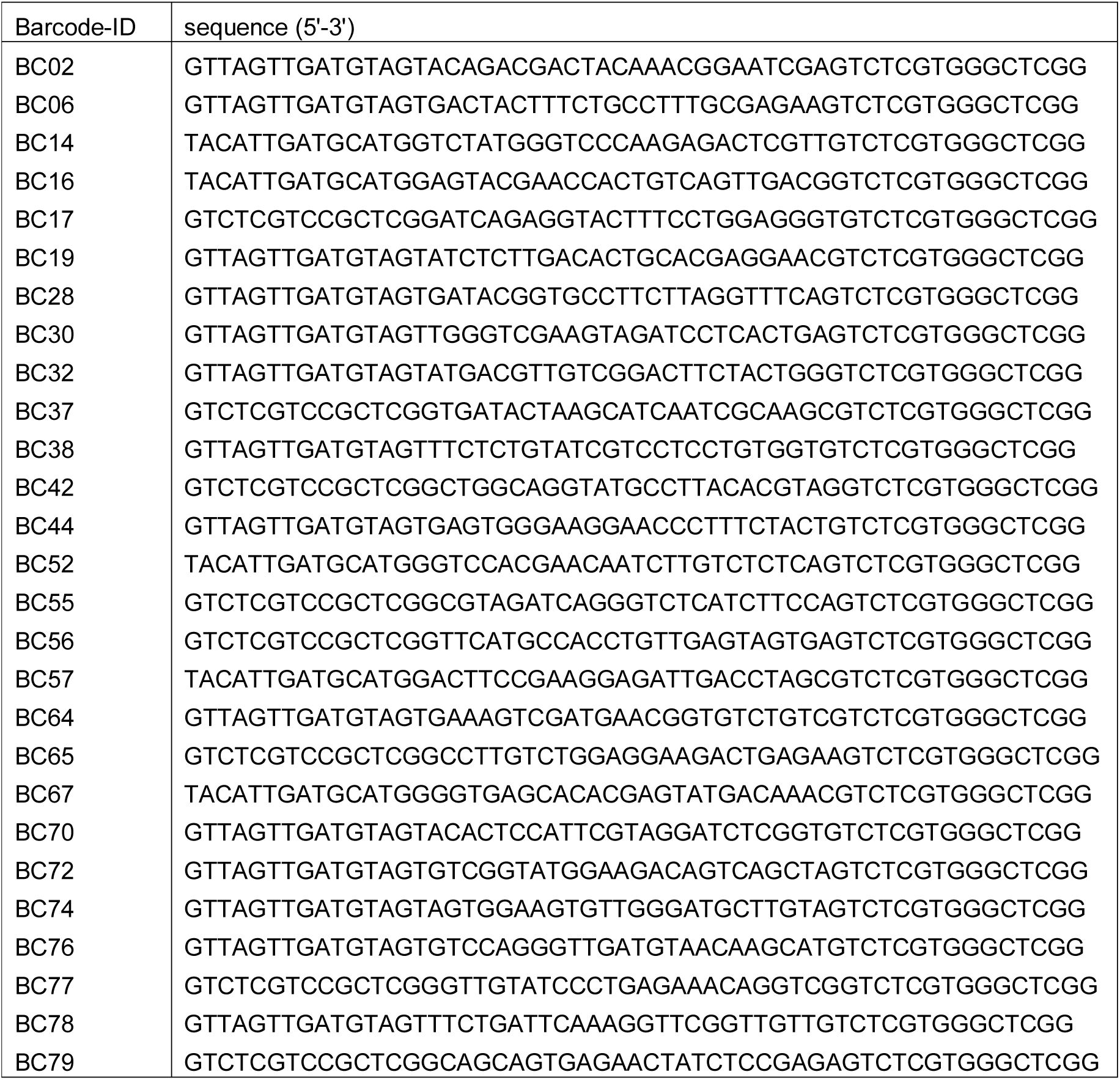
PCR primer sequence for V1-V8 region 16S rRNA gene sequencing with Oxford Nanopore Technologies platform.

**Appendix Fig. 1.**
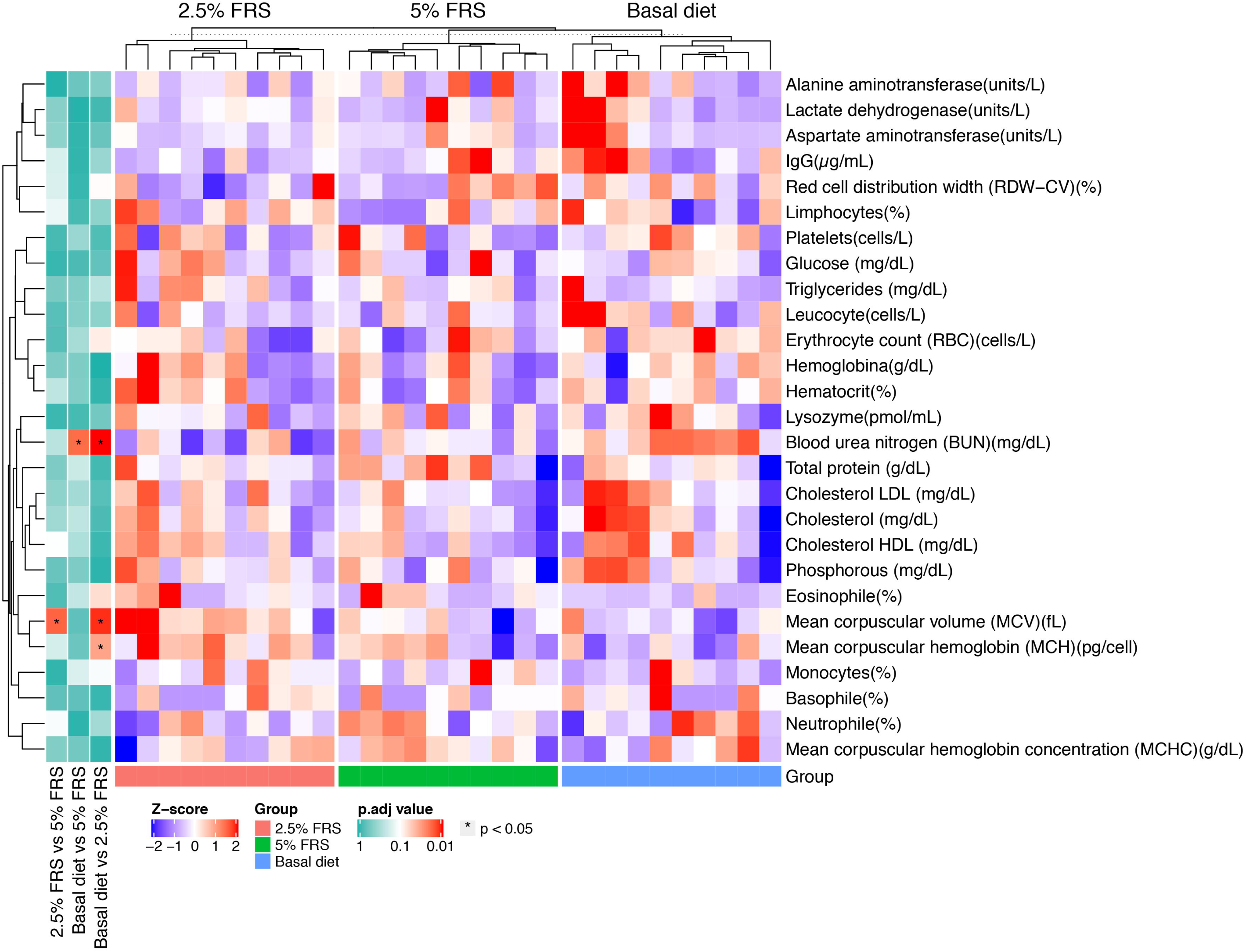
Hierarchical clustered heatmap showing the hematological data of piglets under experimental diets: basal diet, 2.5% and 5% FRS. Respectively n= 10, 10, 10 for basal diet with 0%, 2.5% and 5% added FRS (fermented rapeseed-*Sacharina latissima*-*Ascophillum nodossum*). The *p* values from pairwise comparison is showed by color depth and the labels of * represent *p* < 0.05.

**Appendix Fig. 2.**
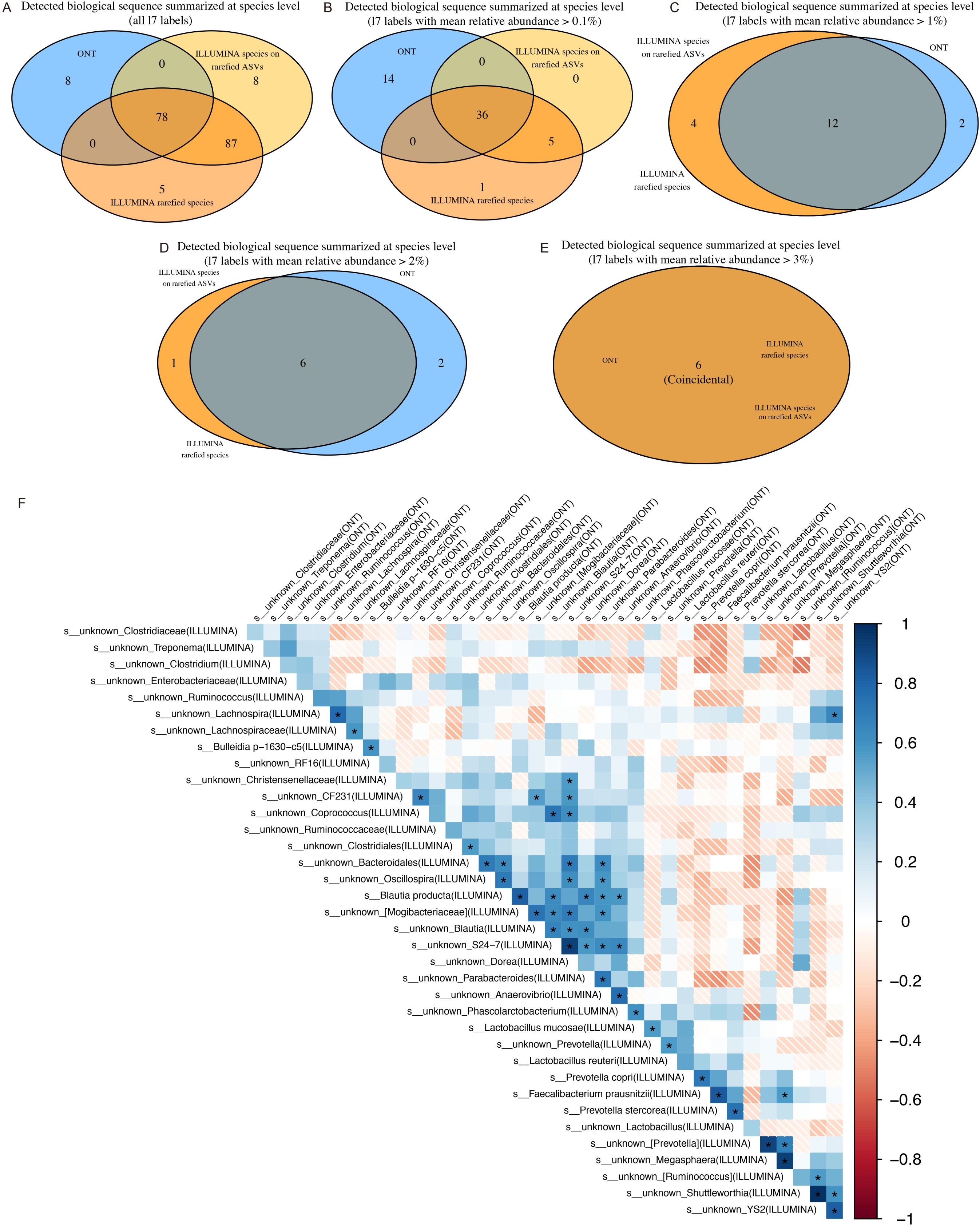
Shared species between the short-read and long-read amplicon sequencing (A-E) and upper triangle heatmap of Pearson’s correlation within the shared taxa (F). Venn plots of species-level summarized features from Illumina and ONT sequencing with mean relative abundance cut-off of 0% (A), 0.1% (B), 1% (C), 2% (D), 3% (E). The annotations of Illumina species on rarefied ASVs, Illumina rarefied species and ONT indicate the captured labels at lowest taxonomic level using the rarefied Illumina ASV table, Illumina and ONT species-level summarized tables respectively.

**Appendix Fig. 3.**
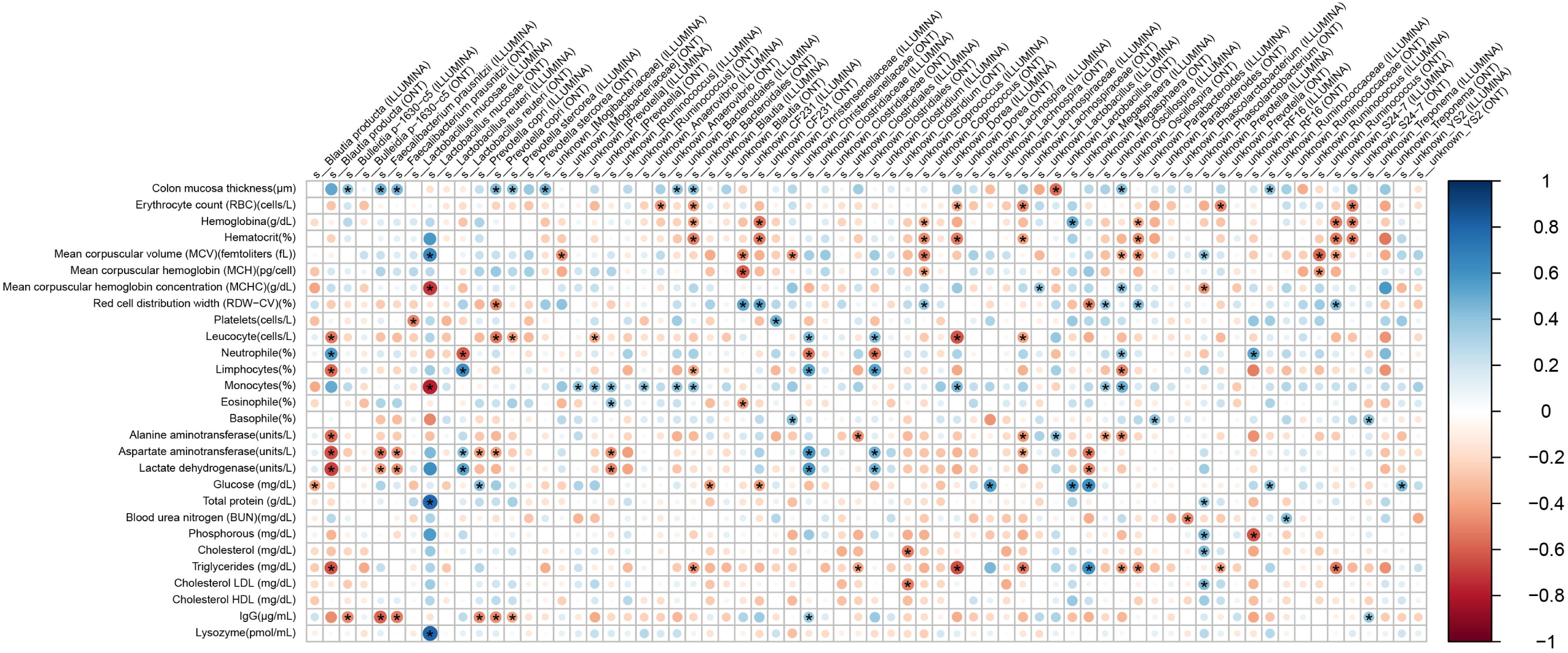
Heatmap showing Pearson’s correlation between the hematological parameters and bacteria relative abundance calculated from V3 region sequencing of 16S rRNA gene and V1-V8 region sequencing of 16S rRNA gene. The color depth and size of the points represent the coefficient and *p* value respectively. Statistically significant pairs with *p* < 0.05 are marked with *.

